# Visualizing and isolating iron-reducing microorganisms at single cell level

**DOI:** 10.1101/2020.09.09.290734

**Authors:** Cuifen Gan, Rongrong Wu, Yeshen Luo, Jianhua Song, Dizhou Luo, Bei Li, Yonggang Yang, Meiying Xu

**Author notes:** Corresponding author: Y. Yang. Guangdong Institute of Microbiology, Guangzhou 510070, China.

## Abstract

Iron-reducing microorganisms (FeRM) play key roles in many natural and engineering processes. Visualizing and isolating FeRM from multispecies samples are essential to understand the in-situ location and geochemical role of FeRM. Here, we visualized FeRM by a “turn-on” Fe^2+^-specific fluorescent chemodosimeter (FSFC) with high sensitivity, selectivity and stability. This FSFC could selectively identify and locate active FeRM from either pure culture, co-culture of different bacteria or sediment-containing samples. Fluorescent intensity of the FSFC could be used as an indicator of Fe^2+^ concentration in bacterial cultures. By integrating FSFC with a single cell sorter, we obtained three FSFC-labeled cells from an enriched consortia and all of them were subsequently evidenced to be capable of iron-reduction and two unlabeled cells were evidenced to have no iron-reducing capability, further confirming the feasibility of the FSFC.

**Importance:** Visualization and isolation of FeRM from samples containing multispecies are commonly needed by researchers from different disciplines, such as environmental microbiology, environmental sciences and geochemistry. However, no available method has been reported. In this study, we provid a solution to visualize FeRM and evaluate their activity even at single cell level. Integrating with single cell sorter, FeRM can also be isolated from samples containing multispecies. This method can be used as a powerful tool to uncover the in-situ or ex-situ role of FeRM and their interactions with ambient microbes or chemicals.

## 1. Introduction

Iron minerals are widespread in anoxic subsurface environments and can be used as electron acceptors by many microorganisms.^1^ In natural environments, those iron-reducing microorganisms (FeRM) not only play a key role in the reduction of minerals and humic substances but also participate in the oxidation of sulfur compounds and organic matters.^2-4^ Moreover, FeRM are important in many engineered processes such as the wastewater treatment, bioremediation and bioelectrochemical systems.^5^ Microbial iron-reducing process is an ancient respiration on the earth.^1^ However, many novel electron transfer strategies (e.g. bacterial nanowire, direct inter-cellular electron transfer) possessed by FeRM were just recognized in recent years.^6,7^

To explore the role of FeRM in various environments, visualizing FeRM is a common need by researches on environmental, microbiological and earth sciences as it can provide essential information such as the location, amount or even activity of FeRM. However, FeRM are phylogenetically rather ubiquitous and thus there is no 16S rRNA or functional gene based assay to detect them so far. Phenanthroline-based spectrophotometric method has been used mostly to evaluate the capability of FeRM.^8^ However, this method is unavailable to identify, locate or quantify FeRM from a multispecies consortia. Recently, some methods targeting cytochromes or extracellular electron transfer processes similar to iron reduction (including azo-dye reduction, tungsten trioxide reduction) have been reported.^9-11^ However, these methods are unsuitable for visualizing FeRMs in a multispecies consortia, because (i) cytochrome proteins are commonly shared by both FeRM and other bacteria; (ii) some FeRM do not reduce other extracellular electron acceptors^12^; (iii) it is hard to use such methods to locate FeRM in complex samples at single cell level.

A common characteristic of FeRM is that, the Fe^2+^ phosphate or carbonate generated from iron reduction can be adsorbed by extracellular polymeric substances, and thus creating a Fe^2+^-accumulated layer on the cell surface of FeRM.^13-15^ This Fe^2+^-layer can be maintained by the reducing forces from outer membrane redox proteins such as *c*-type cytochromes. Therefore, Fe^2+^-selective fluorescent chemosensor may provide a convenient and sensitive tool to visualize FeRM in different environments. Several Fe^2+^-specific fluorescent probes have been developed for mammalian cells but none of them has been tested in microorganisms.^16-19^ In contrast to the intracellular Fe^2+^ detection in mammalian cells, several challenges must be considered for a fluorescent probe for FeRM. For examples, the FeRM-probe should be nonreactive to other microorganisms as Fe^2+^ can also be adsorbed to the surfaces of them, especially in environments containing high concentration of Fe^2+^.

Once FeRM cells can be visualized with fluorescence, single cell sorting techniques (e.g. microfluidic devices, laser tweezer or laser ejection) can be used to isolate them and their partner microbes from where they are observed. Thus, both in situ and ex-situ roles and mechanisms of each targeted FeRM cell can be understood. Guided by this aim, we synthesized an oxygen-depleting Fe^2+^-specific fluorescent chemosensor (FSFC) which showed high sensitivity and selectivity to Fe^2+^. This FSFC also showed high feasibility in visualizing FeRM in pure-cultured and multispecies systems. Integrating with single cell sorting technique, this probe could facilitate identification and isolation of FeRM from an enriched sediment consortia.

## 2. Materials and methods

### 2.1 Synthesis of N-butyl-4-phenyltellanyl-1, 8-naphthalimide (FSFC)

This probe was selected due to its repeatable and simple synthesis method (Fig. S1).^16^ Firstly, 4-bromo-N-butyl-1, 8-naphthalimide was synthesized.^16,20^ In brief, 5.0 g 4-bromo-1,8-naphthalic anhydride and 3 mL n-butylamine were dissolved in 90 mL ethanol and refluxed in 82 °C for 6 h. Then the mixture was filtered to obtain a wine-red solution. After evaporation with a rotary evaporator (90 °C, 80 rpm, until all ethanol was evaporated), the crude product was purified by column chromatography (silica gel, ethyl acetate: petroleum ether = 1: 50) to get a pale yellow solid product (4.8 g). Secondly, N-butyl-4-phenyltellanyl-1, 8-naphthalimide (FSFC) was synthesized by a modified method based on a previous report.^16^ 1.02 g diphenyl ditelluride and 60 mL ethanol were added to a 150 mL three-neck flask flushed with nitrogen. The suspension was cooled to 0 °C, then 0.24 g sodium borohydride was dissolved in 12 mL ethanol and slowly dropt into the three-neck flask. After the red color faded, the reaction mixture was heated to reflux in 83 °C. Then, a mixture of cuprous iodide (0.41 g, 2.2 mmol) and 4-bromo-N-butyl-1, 8-naphthalimide (0.59 g, 1.8 mmol) were added. The mixture was stirred and refluxed for 30 min in a nitrogen atmosphere. After cooling to room temperature, the black mixture was filtered to remove insoluble materials. Then, the black filtrate was evaporized on a rotary evaporator. The residue was washed by ethanol and filtered again. After evaporation (90 °C, 80 rpm, until all ethanol was evaporated), the residue was purified by column chromatography (silica gel, ethyl acetate: petroleum ether = 1:125) to obtain a yellow solid product of FSFC (0.72 g). The final yellow product was dissolved in acetonitrile to get a 5 mM stock solution. It was further diluted by phosphate buffer saline (PBS, containing 3.6 g/L Na_2_HPO_4_·7H_2_O, 0.27 g/L KH_2_PO_4_, 8 g/L NaCl, and 0.2 g/L KCl, pH 7.2) and stored in dark before use.

### 2.2 Sensitivity and selectivity test of FSFC

Aqueous solutions of ferric citrate (FeC_6_H_5_O_7_·5H_2_O), ammonium iron (II) sulfate hexahydrate (H_8_FeN_2_O_8_S_2_·6H_2_O), MnCl_2_, ZnCl_2_, CaCl_2_, MgCl_2_, NiCl_2_·6H_2_O, CuCl_2_, Co(NO_3_)_2_·6H_2_O, CdCl_2_·2.5H_2_O, NaCl and KCl, were used for the selectivity and sensitivity tests of Fe^3+^, Fe^2+^, Mn^2+^, Zn^2+^, Ca^2+^, Mg^2+^, Ni^2+^, Cu^2+^, Co^2+^, Cd^2+^, Na^+^, K^+^ respectively. Millipore water was used to prepare all kinds of aqueous solution. For each experiment, fresh Fe^2+^ solution was prepared before using.

Sensitivity of FSFC towards Fe^2+^ was tested by a PerkinElmer LS 45 fluorescence spectrometer. Typically, the sensitivity of FSFC was carried by incubating the FSFC (50 μM) with 0, 5 µM, 10 µM, 20 µM, 50 µM, 100 µM, 200 µM, 500 µM, 1000 µM and 2000 µM of Fe^2+^ (ammonium iron sulfate hexahydrate, H_8_FeN_2_O_8_S_2_·6H_2_O) for 30 min. The reaction solution (3 mL final volume for each solution) was added into a quartz cell for fluorescence measurements with an excitation wavelength (λ_*ex*_) at 445 nm and the emission wavelength (λ_*em*_) from 480 nm to 600 nm.

Selectivity of FSFC towards Fe^2+^ was investigated by incubating 50 μM FSFC with 100 μM of various specified cations (Fe^3+^, Fe^2+^, Mn^2+^, Zn^2+^, Ca^2+^, Mg^2+^, Ni^2+^, Cu^2+^, Co^2+^, Cd^2+^, Na^+^, K^+^ in ferric citrate (FeC_6_H_5_O_7_·5H_2_O), ammonium iron (II) sulfate hexahydrate (H_8_FeN_2_O_8_S_2_·6H_2_O), MnCl_2_, ZnCl_2_, CaCl_2_, MgCl_2_, NiCl_2_·6H_2_O, CuCl_2_, Co(NO_3_)_2_·6H_2_O, CdCl_2_·2.5H_2_O, NaCl and KCl), respectively for 30 min. The reaction solution (3 mL final volume for each solution) was added into a quartz cell for fluorescence measurements with an excitation wavelength (λ_*ex*_) at 445 nm and the emission wavelength (λ_*em*_) at 530 nm.

### 2.3 Pure culture strains and growth conditions

To test the selectivity of FSFC to FeRM, pure bacteria cultures including three known FeRM (*Shewanella decolorationis* strain S12, *S. oneidensis* MR-1 and *Geobacter sulfurreducens* PCA),^5,6,21,22^ three non-FeRM (a *ccmA*-mutant *S. decolorationis* S22, *Massilia rivuli* FT92W, *Duganella lacteal* FT50W) incapable of iron-reduction,^21,23^ and five pure cultured bacteria newly isolated from sediments with unknown iron-reduction capacity (*Paenibacillus motobuensis* Iβ12, *Ciceribacter* sp. F217, *Sphingobium hydrophobicum* C1, *Bacillus* Iβ8, *Lysinibacillus varians* GY32) were used. Further information of those bacteria were listed in Table S1. All bacteria (except for *G. sulfurreducens* PCA) were firstly grown aerobically in Luria-Bertani (LB) medium. The bacteria cells were washed with sterilized PBS for two times and then inoculated to N_2_-flushed anaerobic lactate medium (LM, containing 2.0 g/L lactate, 0.2 g/L yeast extract, 12.8 g/L Na_2_HPO_4_·7H_2_O, 3 g/L KH_2_PO_4_, 0.5 g/L NaCl, and 1.0 g/L NH_4_Cl) with an initial OD_600_ of 0.1. 3 mM of ferric citrate or Fe_2_O_3_ was used as the electron acceptor in the LM, unless otherwise stated. The cultures were grown at 33 °C. *G. sulfurreducens* PCA (initial OD_600_ = 0.08) was anaerobically cultivated using freshwater medium containing acetate (10 mM) as electron donor and ferric citrate (4 mM) or poorly crystalline Fe(III) oxides (4 mM) as electron acceptor.^6^ The poorly crystalline Fe(III) oxides were prepared as previous report.^6^ Meanwhile, the iron reducing capability of those bacteria was tested with traditional Phenanthroline-based method.^8^

### 2.4 Co-culture grown in liquid medium and sediment

Different co-culture systems was used to test whether FSFC can distinguish the FeRM from non-FeRM in the same bacteria culture. The co-culture systems includes: (i) co-culture system of *S. decolorationis* S12 and non-FeRM *L. varians* GY32 in N_2_-flushed anaerobic LM solution with 3 mM of ferric citrate for 8 h; (ii) 10 mL of the above co-culture system was inoculated with 1 g of sterilized river sediment (obtained from Shijing River, Guangzhou, China), cultivated for 8 h.

### 2.5 Fluorescence Imaging

The fluorescence imaging of the pure culture or co-culture samples were obtained with a confocal laser scanning microscopy (CLSM, LSM 700, Zeiss) after being stained by 50 μM FSFC for 15 min. FSFC concentration higher than 0.5 mM may cause toxicity to bacteria (Fig. S2). 10 μL of the stained cultures was dripped on a glass slide with small piece of cover slide and then observed under the CLSM with an excitation wavelength (λ_ex_) at 445 nm for FSFC. Propidium iodide (PI, λ_ex_= 490 nm, Thermo Fisher) was used as a fluorescent indicator for evaluating the activity of the bacterial cells, only cells with low activities and impaired cell membrane can be stained by PI.

### 2.6 FSFC-based single cell isolation

An enriched iron-reducing biofilm consortia was used to test that whether FSFC can selectively label FeRM in a complex microbial community. This consortia was made by inoculating 1 g of sediment into 100 mL LM containing 5 mM of ferric citrate in an anaerobic serum bottle. Six graphite plates (1 × 1 × 0.1 cm) were added in the culture for biofilm growth. 80% of the enriched culture was replaced with fresh LM containing 5 mM of ferric citrate for every two weeks. After being enriched for eight-weeks, three of the graphite plates were fetched and stained by FSFC and PI after a gentle wash in sterilized PBS. The stained biofilms were observed under CLSM. Biofilms on the other three graphite plates were scraped by a sterilized cotton swab. The resulting biofilm cells were suspended in 5 ml PBS and stained by 50 μM FSFC for 15 min. 50 μL of the FSFC-stained sample was transfer to a glass slide designed for single cell ejection and observed under fluorescence mode of the single cell precision sorter (PRECI SCS, HOOKE Instruments). The selected cells (with or without fluorescence) on the slides were ejected by a laser beam controlled by PRECI SCS software. 7 bacterial cell with fluorescence and 6 bacterial cells without fluorescence were ejected from the slide into a collector containing sterilized PBS by low power laser (0.5 to 1 μJ, varied according to the cell shape and adsorption force on the slide surface). The collected single bacteria was then anaerobically cultivated in LB medium containing 2 mM of ferric citrate. The grown bacteria were then cultivated in the same freshwater medium used for *G. sulfurreducens* PCA with ferric citrate as sole electron acceptor and acetate as electron donor.

## 3. Results and discussions

### 3.1 Sensitivity, selectivity and stability of FSFC

FSFC was non-fluorescent (“off” state) in the absence of Fe^2+^ due to the heavy-atom effect of the tellurium atom on the naphthalimide fluorophore. Fe^2+^ can trigger the detelluration reaction of FSFC and cause a strong fluorescence (“on” state).^16^ As evidenced by GC-MS (Fig. S3), the purity of the FSFC product was 94.2%. Fig. 1A showed that FSFC exhibit very weak background fluorescence in the absence of Fe^2+^. Upon the addition of Fe^2+^ from 0 to 2000 μM, the fluorescence emission increased accordingly. Fig. 1B showed a linear relation between the fluorescence intensity (FI) and the logarithm of Fe^2+^ concentration. The theoretical limit of detection (LOD) was calculated to be 6.3 μM (based on the formula LOD = 3 × σ/m, σ is the standard deviation of the response at the lowest tested concentration and m is the slope of the concentration-FI response).^24^ Generally, the concentration of Fe^2+^ in practical environments varied from several to hundreds of μM and could be up to several mM in FeRM cultures.^25,26^ Therefore, FSFC could be used as an alternative Fe^2+^ sensor or FeRM-label for most environmental and experimental samples.

**Fig. 1.**
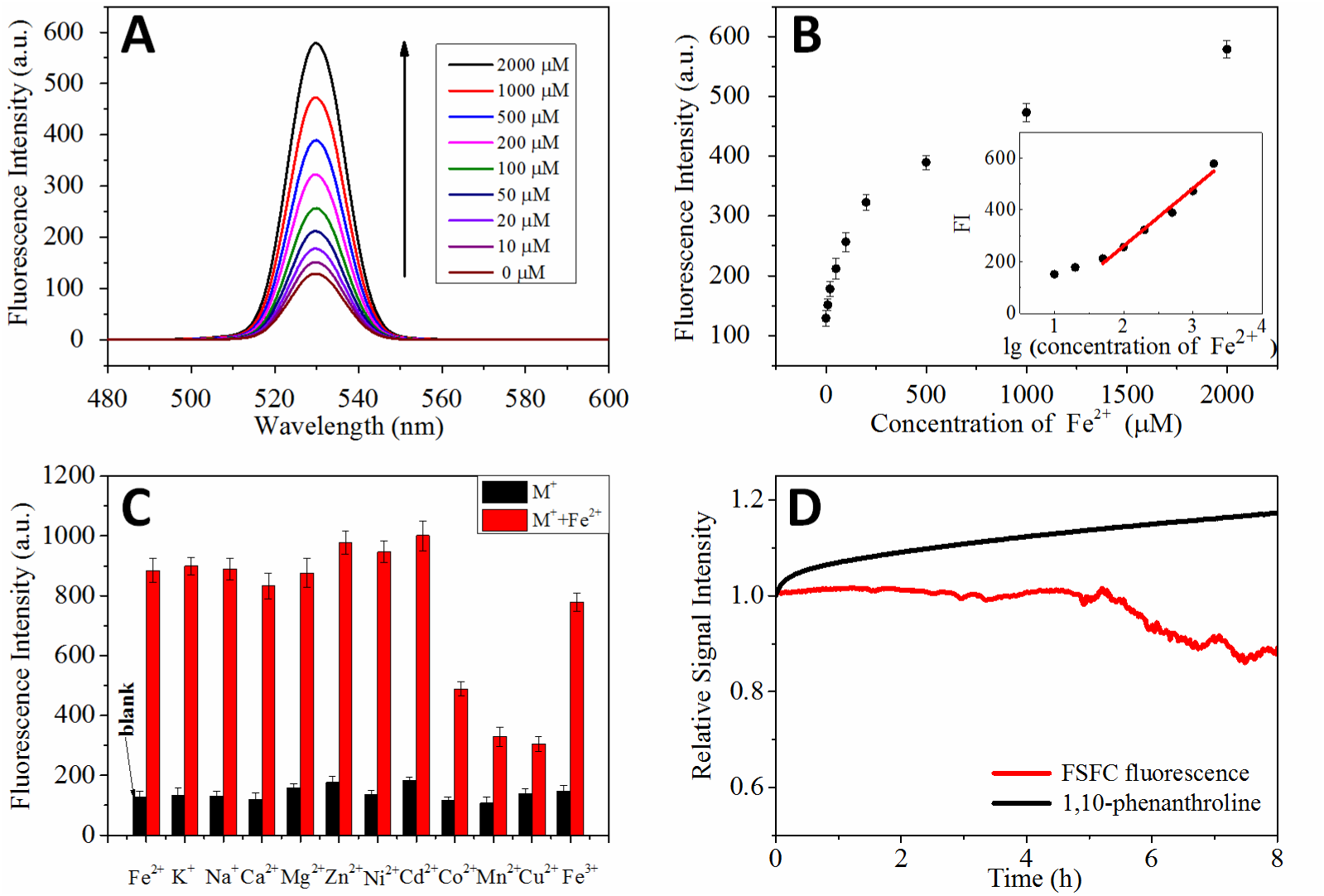
The sensitivity, selectivity and stability of FSFC in Fe^2+^-containing solution. (A) Response of FSFC fluorescence spectra to different concentrations of Fe^2+^. (B) Relationship between the concentration of Fe^2+^ and the FI. Insert shows the linear relationship between FI and the logarithm of Fe^2+^ concentrations. (C) Selectivity tests of FSFC to Fe^2+^. Black bars indicate fluorescence response of FSFC to deionized water (blank) and deionized water containing different metal cations (M^+^), red bars indicate fluorescence response of FSFC to different cations combined with Fe^2+^. (D) Relative stability of FSFC and traditional o-phenanthroline-based method.

Other metal ions in practical environments may affect the fluorescence response of FSFC to Fe^2+^. Fig. 1C showed that all tested metal ions (except for Fe^2+^) had no significant fluorescence response to FSFC individually. When co-existing with Fe^2+^, metal ions K^+^, Na^+^, Ca^2+^, Mg^2+^, Zn^2+^ had little effects on the Fe^2+^-FSFC fluorescence while Cu^2+^, Mn^2+^ and Co^2+^ could affected the fluorescence to some extents. In typical natural environments, the concentrations of Co^2+^, Cu^2+^ and Mn^2+^ are generally several orders of magnitude lower than that of Fe^2+^,^25^ e.g. the concentration of manganese was two orders of magnitude lower than that of iron (8 vs 800 μM) in the sediments of Yaquina Bay Estuary,^25^ indicating that the effects of other metal ions will be small for tests with naturally environmental samples. It should be noted the effects of other metal ions on FSFC fluorescence may increase with their concentrations (Fig. S4). For some industrial wastewaters containing high concentration of metal ions, the samples should be diluted or Fe^2+^ should be artificially elevated before using FSFC to visualize FeRM.

Fig. 1D showes a stability comparison between FSFC fluorescence and the traditional o-phenanthroline method. The FI of FSFC remained stable within 5 h (deviation < 5%) while the signal of traditional phenanthroline-method increased by over 10% within 2 h. Therefore, FSFC had better stability (within 5 h) compared to the phenanthroline-method, which also means that FSFC has particularly advantage in the studies needing long-time operations or including large number of samples.

### 3.2 Fluorescence imaging of viable FeRM reducing soluble and solid Fe^3+^

Iron reducing capability of *Shewanella* and *Geobacter* species has been extensively demonstrated in previous studies^2,6,7,15,27,28,^. Moreover, it has been reported that Fe^2+^ phosphate and carbonate aggregate on cellular surfaces during the iron-reduction by FeRM.^15^ Our results also showed that compared to the non-FeRM (0.1-0.3 μM/μg bacteria protein),much higher concentration of Fe^2+^ accumulated on the cell surface of *S. decolorationis* S12 (1.1 μM/μg bacteria protein) and *S. oneidensis* MR-1 (1.2 μM/μg bacterial protein) when exposing to the same Fe^2+^-containing culture (Fig. S5), which further supported the reasonability of using Fe^2+^-probe to identify FeRM. Fig. 2 showed that the *S. decolorationis* S12 in iron-reducing medium displayed significant fluorescence while the cells grown aerobically (without Fe^3+^) have no fluorescence, indicating that cell surface-adsorbed Fe^2+^ can selectively turn-on the fluorescence of FSFC. Moreover, the FI on S12 cell surface increased correspondingly with the Fe^2+^ concentration in the culture supernatant (Fig. 2B-D). By integrating with PI, a fluorescent dye targeting inactive bacteria (with impaired cellular membrane), it can be seen that FSFC only label the active iron-reducing S12 cells (Fig. S6). This result demonstrated that FSFC was selectively targeted to the active iron-reducing strain S12 cells rather than the inactive or non-FeRM strain S12 cells, probably because the Fe^3+^-reducing activity of inactive cells was low and thus the Fe^2+^ accumulation layer cannot be maintained on the surfaces of such cells.

**Fig. 2.**
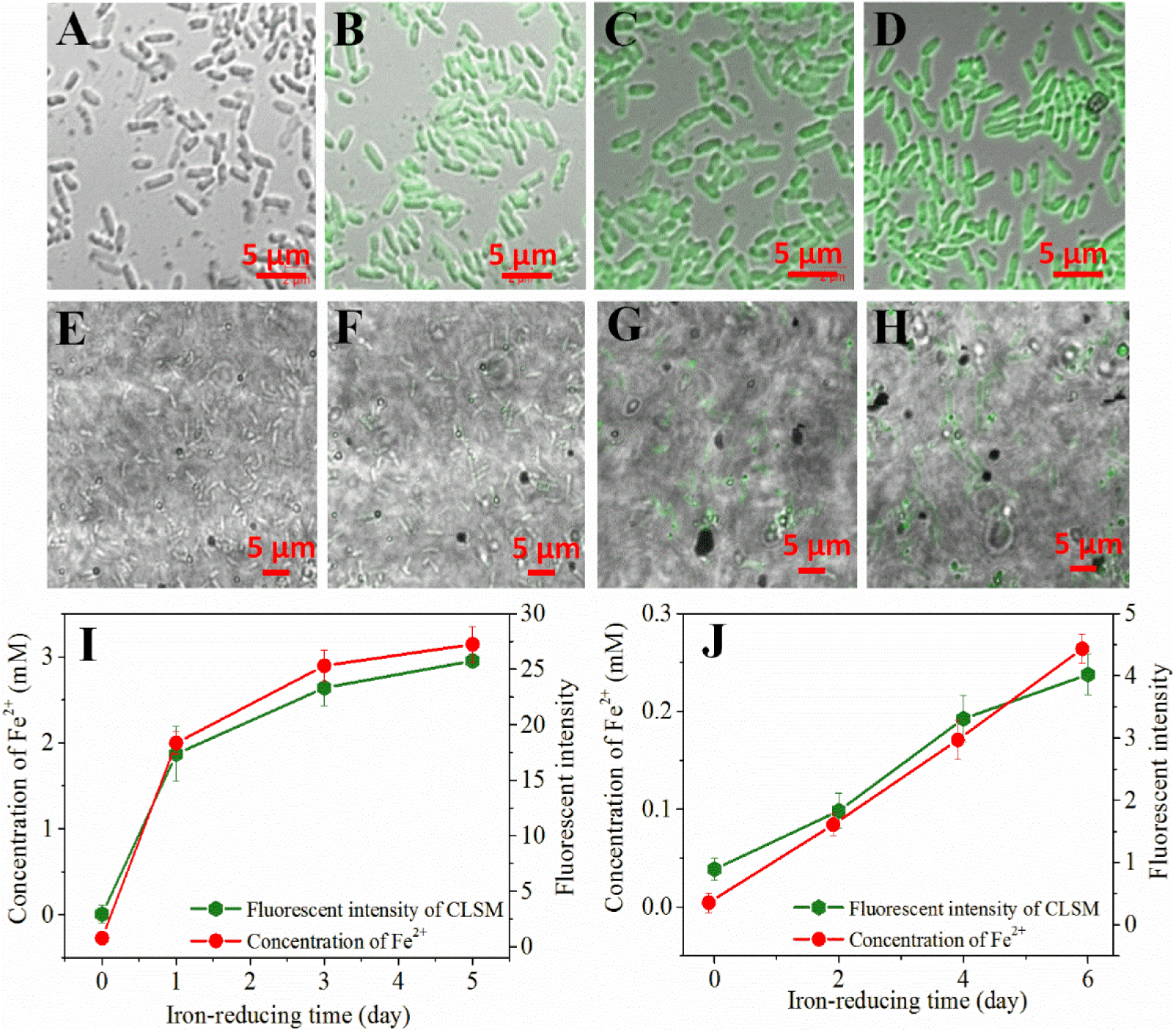
Fluorescence response of FSFC to strain S12 using oxygen, soluble Fe^3+^ or solid Fe^3+^ as electron acceptor. (A) S12 respiring with oxygen or at 0 h in Fe^3+^-reducing medium; (B-D) S12 respiring with soluble Fe^3+^ for 1, 3, 5 h, respectively; (E-H) S12 respiring with solid Fe^3+^ for 0, 3, 5, 7 days, respectively; (I, J) Fe^2+^ concentration and the corresponding FI of strain S12 with soluble Fe^3+^ or solid Fe^3+^, respectively.

Considering that iron exist mainly as solids in natural environments, the reduction process of Fe_2_O_3_ particles by strain S12 was also investigated. Due to the low reducing capability of strain S12 on Fe_2_O_3_, almost no fluorescence was observed in the first 2 days (Fig. 2E, F). The results showed that FSFC had no fluorescence response to Fe_2_O_3_ particles. Over the next 5 days, fluorescence on S12-cells gradually increased with the increase in Fe^2+^ concentration (Fig. 2G-H). The FI was much lower of S12 grown with Fe_2_O_3_ particles compared to that with soluble Fe^3+^, which was corresponding to the different reduction rates of strain S12 with the two forms of Fe^3+^. In the system with either soluble or solid Fe^3+^, FI on the cells showed linear relationship to the ambient Fe^2+^ concentration (Fig. 2I, J). We also tested the performance of FSFC with *S. oneidensis* MR-1 redcuing soluble or solid Fe^3+^ which showed consistent results with that with *S. decolorationis* S12 (Fig. S7).

*Geobacter* has different extracellular electron transfer pathways compared to *Shewanella*.^5,7^ When using ferric citrate as electron acceptor, the FI on *G. sulfurreducens* PCA cell increased with the Fe^2+^ concentration which was similar to the two *Shewanella* species. However, when reducing solid Fe^3+^ oxides, only *G. sulfurreducens* PCA cells attached to the Fe^3+^ particles showed fluorescence while planktonic cells showed weak or no fluorescence (Fig. S8). The different fluorescent performances between *Shewanella* and *Geobacter* when reducing solid Fe^3+^ electron acceptors may be explained by their extracellular electron transfer pathways: *Shewanella* can secret soluble electron mediators to dissolve and reduce Fe^3+^ particles without attaching to the particles while *G. sulfurreducens* PCA can only reduce Fe^3+^ particles via outer membrane cytochrome *c* or e-pili after attaching to the particles.^5,6,7^ These results demonstrated that FSFC can visualize FeRM reducing either soluble or solid Fe^3+^. Moreover, the FI on the bacteria surface can be considered as an indicator of the Fe^2+^ concentration in the pure cultures reducing soluble Fe^3+^. However, it should be noted that the FI of different *G. sulfurreducens* PCA cells on the same Fe^3+^ aggregates varied largely (Fig. S8), indicating different physiological status of them at single cell level.

### 3.3 Evaluating the iron-reducing capability of different bacteria

In addition to iron-reducing capability, bacteria from different genera usually have different shapes, surface properties and metabolites that may affect the fluorescence of FSFC. To further analyze the selectivity of FSFC, we used FSFC to test five blind bacterial samples (five bacteria newly isolated from sediment with unknown iron-reducing performance, Table S1), with *S. decolorationis* S12, *S. oneidensis* MR-1 as positive controls (capable of iron-reduction) and *ccmA*-mutant S22 (deficiency in producing mature c-type cytochromes),^21^ *Massilia rivuli* FT92W, *Duganella lacteal* FT50W as negative controls (incapable of iron-reduction).^23^ As expected, *S. decolorationis* S12, *S. oneidensis* MR-1 showed fluorescence while the negative controls showed no fluorescence (Fig. 3, Fig. S9). Among the five blind bacterial samples, only *Paenibacillus motobuensis* Iβ12 had fluorescence but the FI was lower than that of *S. decolorationis* S12. The other bacteria have no fluorescence (Fig. 3A-G). Traditional o-phenanthroline method showed consistent results that only *P. motobuensis* Iβ12 had iron-reducing capability and its iron-reducing rate is much lower than that of *S. decolorationis* S12 (0.14 vs 0.58 mM/h). The results indicated that FSFC could be used as a simple and visualizing method to identify and evaluate the iron-reducing capability of different bacteria.

**Fig. 3.**
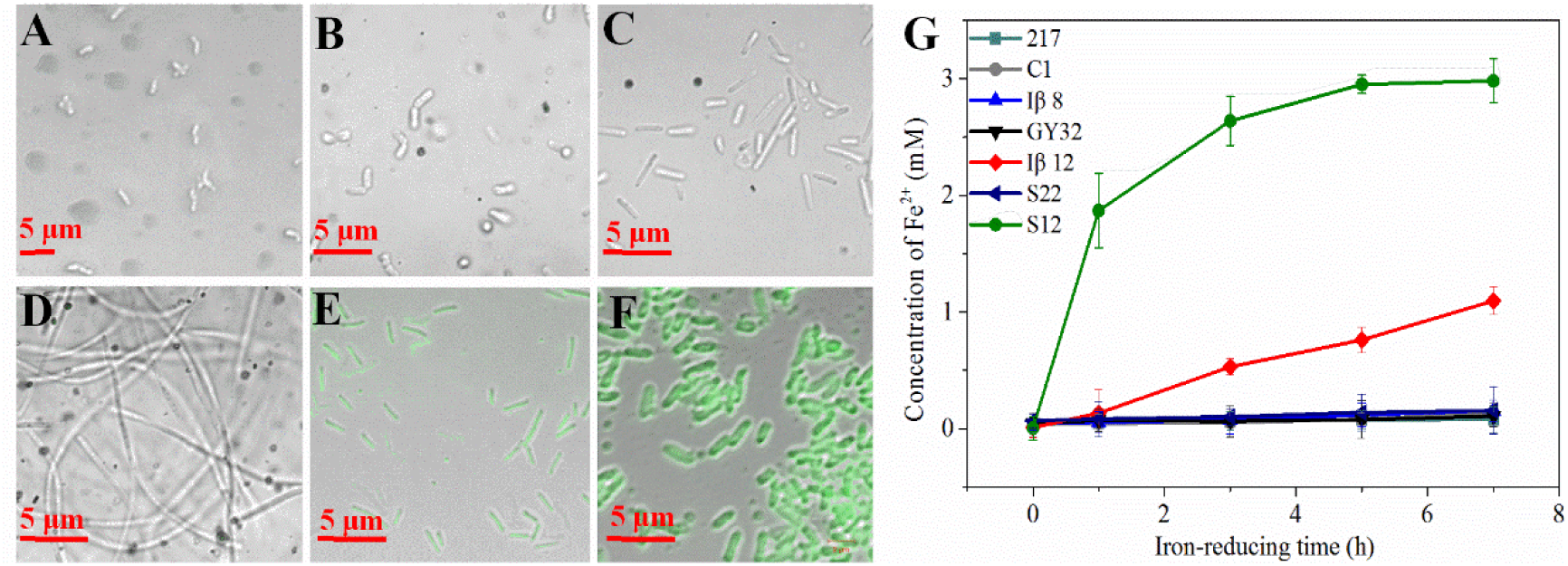
Fluorescence images of FSFC to different bacterial cultures containing ferric citrate. (A) Ciceribacter sp. F217, (B) S. hydrophobicum C1, (C) Bacillus Iβ8, (D) L. varians GY32, (E) P. motobuensis Iβ12, (F) S. decolorationis S12, (G) the iron-reduction of different strains. (Scale bar: 5 μm)

### 3.4 Visualizing FeRM from bacterial co-cultures

Co-culture of FeRM and bacteria with other functions is an important way to understand the interaction between FeRM and other bacteria. In such co-culture systems, one possible problem challenging FSFC is that the Fe(II) generated by FeRM may adsorbed to non-FeRM and render the later fluorescence. To test whether FSFC can identify FeRM in co-culture systems, we co-cultured a filamentous non-FeRM *L. varians* GY32 and *S. decolorationis* S12 using lactate as electron donor. As shown in Fig. 4A, the rod-shape strain S12 showed strong fluorescence while the filamentous bacteria *L. varians* GY32 have no fluorescence in the same iron-reducing culture. It can be seen that FSFC can selectively visualize the FeRM in microbial samples containing FeRM and non-FeRM. The result was consistent with that Fe^2+^ accumulated on the surface of non-FeRM is much less even in the same Fe^2+^-containing environment (Fig. S5). By integrating with a flow cytometer, we could separate the iron reducing bacterium *S. decolorationis* S12 from a co-culture of two rod-shape bacteria (*S. decolorationis* S12 and *S. hydrophobicum* C1, Fig. S11) by the fluorescence, suggesting potential application of FSFC for FeRM with properly controlled flow cytometer or other microfluidic techniques. However, the bacteria samples for microfluid- or microdroplet-based techniques must be simple and well-separated. The pretreatment of most environmental samples which contain aggregates or filamentous bacteria will be challenging for such microfluidic techniques.

**Fig. 4.**
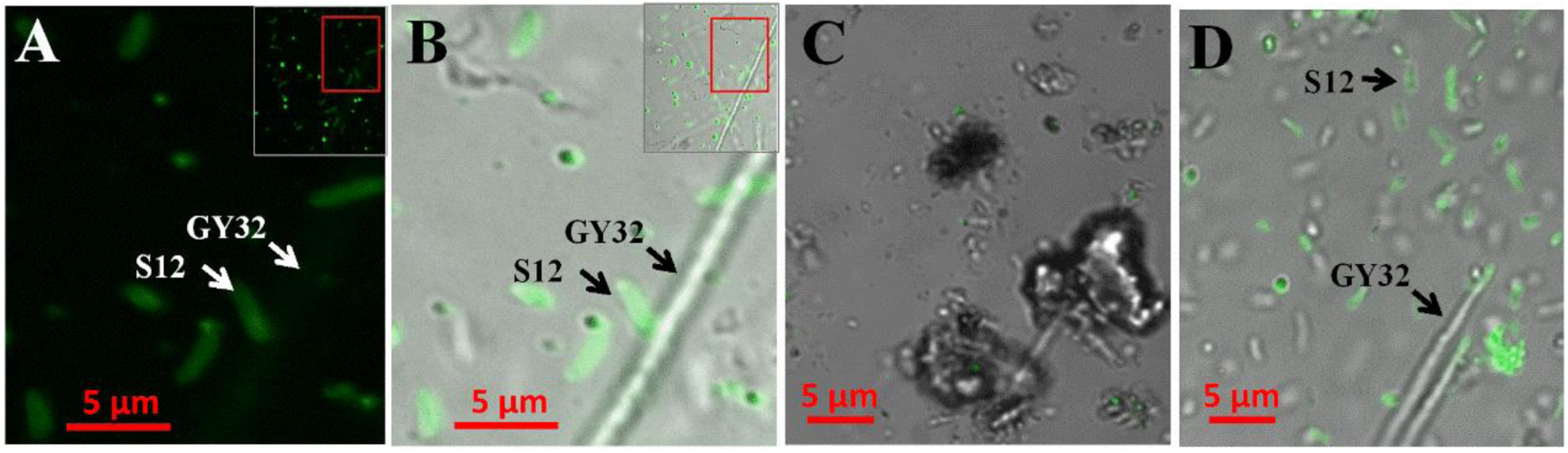
Fluorescence images of S. decolorationis S12 and L. varians GY32 co-culture. (A) Fluorescence mode image of the co-culture (Fe^2+^concentration: 2.3 mM), magnified from the red rectangle area in the insert; (B) Light-fluorescence merged image of the co-culture in liquid medium (Fe^2+^concentration: 2.3 mM), magnified from the red rectangle area in the insert; (C, D) Light-fluorescence merged image of the sediments with and without co-culture, respectively (Fe^2+^concentration: 1.9 mM).

To evaluate the feasibility of FSFC in more complex environments, FSFC was used to the co-culture of *L. varians* GY32 and *S. decolorationis* S12 in sterilized sediment containing ferric citrate. Fig. 4C showed that in the sediments without co-culture, only a minority of particles showed fluorescence probably due to the inherent Fe^2+^ on those sediment particles and no bacteria-like particles showed fluorescence. The results showed that FSFC had little background fluorescence in sediments and the unviable (sterilized) microorganisms in sediment could not trigger the fluorescence of FSFC. In the co-culture system, short-rod strain S12 showed significant fluorescence while the filamentous bacteria *L. varians* GY32 had no fluorescent, indicating the feasibility of FSFC for visualizing FeRM in sediment-containing environments. However, it should be noted that a minor portion of particles in sediments also had fluorescent response to FSFC probably because some particles can absorb the Fe^2+^ generated by FeRM. A proper dilution or filter could be used to remove the particles from the sediment samples.

### 3.5 Visualizing and isolating single cell FeRM from multispecies consortia

In addition to visualizing FeRM, isolating FeRM from multispecies samples is a general and important need for understanding the iron-associated biogeochemical processes.^29^ The selective fluorescent of FSFC to FeRM provide the possibility of isolating single FeRM cell from multispecies with a single cell isolating platform. Fig. S12 shows that *S. decolorationis* S12 can be distinguished and isolated from the co-culture containing wild strain S12 and mutant strain S22 by integrating FSFC with a laser-based single cell sorter. The laser power used to eject the single microbial cell (< 1 μJ) with this platform was three-order of magnitude lower than the power that (several mJ) may hurt cell viability.^30^

We combined FSFC and PI to label the biofilms in an enriched iron-reducing reactor. CLSM showed that the active FeRM cells were mainly located at the outer layer of the biofilms while the inner (bottom) layer biofilms cell showed low activity and little FSFC fluorescence (Fig. 5A). This activity profile was similar with that of the biofilms respiring with nitrate or azo dyes as electron acceptors^31^, indicating the Fe^3+^ was inaccessible to the inner biofilm layers and thus only the outer layer biofilm cells can reduce Fe^3+^ and maintain high activity. Microbial community analysis showed that the diversity of the enriched biofilm consortia was significantly decreased compared to the initial community (Fig. S13). Gram-positive bacteria were dominant in the enriched consortia. After addition of FSFC to the suspended biofilm consortia, both fluorescent bacteria and non-fluorescent bacteria were observed (Fig. S13). Seven single cells with fluorescence and six single cells without fluorescence were isolated from the enriched consortia using the single cell sorter (Fig. 5). Three of the isolated fluorescent single cells (named S1, S2, S3) were successfully cultivated and all of them could use acetate as electron donor to reduce ferric citrate (Fig. 5F). The 16S rRNA genes of the isolated FeRM S1 (accession number MT947627), S2 (accession number MT947628) were close to *L. fusiformis* NBRC15717 (similarity 99.84%) and *L. pakistanensis* NCCP-54 (similarity 100%), respectively. S3 (accession number MT947629) was close to *Paenibacillus glucanolyticus* NBRC 15330 (similarity 99.25%), respectively. *Lysinibacillus* commonly exists in various environments such as sediment or wastewater.^32-34^ Although the capability of several *Lysinibacillus* strains using electrodes as electron acceptors have been reported,^33,34^ our results present the first evidence that the genus *Lysinibacillus* can reduce iron. *Paenibacillus* is also a common gram-positive bacterial genus and several species in this genus have been demonstrated to reduce iron.^35,36^ On the other hand, two of the non-fluorescent single cells with 16S rRNA genes similar to *Bacillus terrae* RA99 (similarity 99.28%, accession number MT947630) and *P. barengoltzii* NBRC 101215 (similarity 99.58%, accession number MT947631) were successfully cultivated and had no iron-reducing capacity (Fig. 5F). *B. terrae* was identified as a new aerobic species from rhizosphere soils while *P. barengoltzii* NBRC 101215 was an aerobic bacterium that can degrade chitin.^37, 38^ The results also suggested that FSFC could be used as a novel and efficient method to isolate FeRM from different environments by integrating with single cell isolation techniques.

**Fig. 5.**
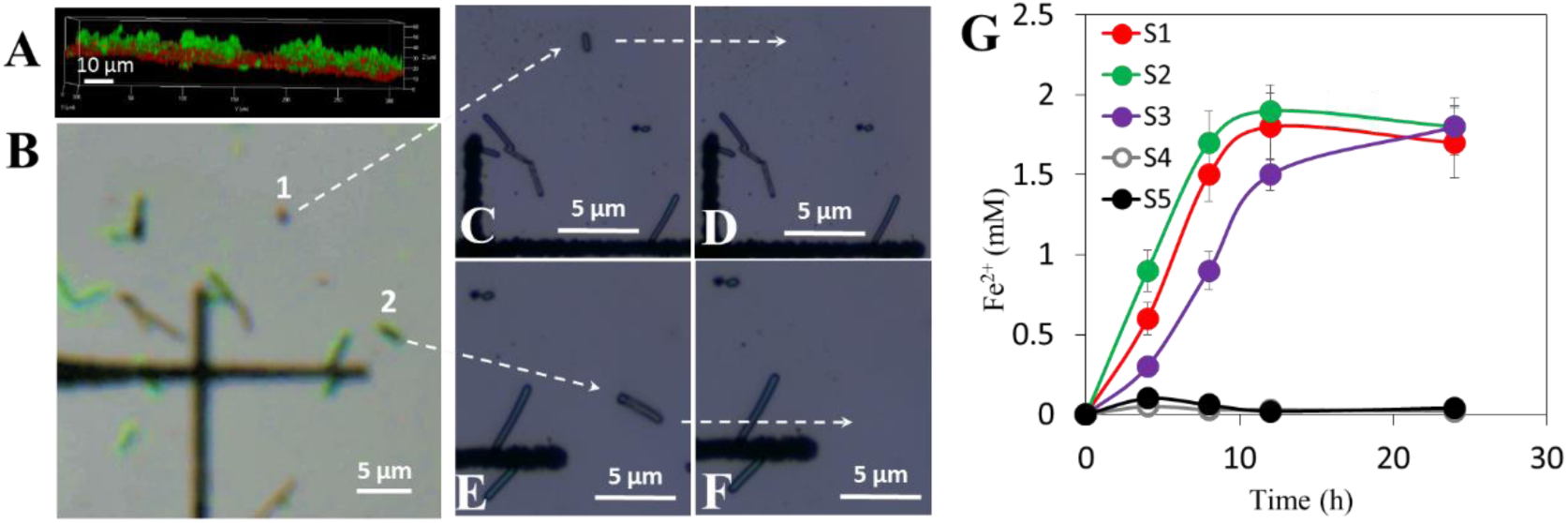
FSFC-based single cell isolation and iron-reducing capability test. (A) Vertical section view of enriched iron-reducing biofilm, red indicates PI-stained cells and green indicates FSFC-labeled cells. (B)Light-fluorescence merged image area of the suspended iron-reducing biofilms. Cell 1 (non-fluorescent) and 2 (fluorescent) are two typically targeted cells to be isolated. The dark cross is a land-mark designed on the glass slide. (C, D) Images before and after the laser-ejection of cell 1 from the slide to a collecting pore containing PBS. (E, F) Images before and after the laser-ejection of cell 2, respectively. (G) Iron-reduction capability of the isolated bacteria.

This study reports a method that can visualize and isolate FeRM from bacterial cultures containing multispecies or even sediments. The FSFC has high sensitivity, selectivity and stability to Fe^2+^ and low background fluorescence in both liquid and sediment environments. In pure cultures or co-cultures containing FeRM, FSFC could selectively visualize the active FeRM. By integrating with single cell sorting technique, targeted FeRM could be efficiently obtained from samples at single cell level. This novel method could be a powerful tool serving for obtaining novel FeRM and for a deeper understanding of the biogeochemical role of FeRM in different environments.

## ASSOCIATED CONTENT

### Supporting Information

Table S1. Information of bacterial strains used in this study.

Figure S1. Synthesis route and function mechanism of N-butyl-4-phenyltellanyl-1, 8-naphthalimide (FSFC).

Figure S2. Toxicity of FSFC on *S. decoloratIonis* S12 cultivated aerobically in LB medium.

Figure S3. GC-MS analysis of the synthesized products.

Figure S4. FI of FSFC in solutions containing 100 μM Fe^2+^ and different concentrations of Mn^2+^.

Figure S5. Fe^2+^ collected from the cell surfaces of different bacteria.

Figure S6. PI-FSFG co-staining on strain S12.

Figure S7. FSFC-stained *S. oneidensis* MR-1 reducing soluble and solid Fe^3+^.

Figure S8. FSFC-stained *G. sulfurreducens* PCA reducing soluble and solid Fe^3+^.

Figure S9. Fluorescent images of control bacteria.

Figure S10. Fluorescent images of S. oneidensis MR-1 when (A) actively reduces ferric citrate and generates 0.1 mM Fe(II) and (B) exposed to 0.1 mM dissolved Fe2+.

Figure S11. Flow cytometry scatter plots of strain S12 and *S. hydrophobicum* C1.

Figure S12. Single cell isolation and determination of strain S12 and *ccmA*-mutant S22 from their co-culture.

Figure S13. Microbial composition and the FSFC-staining of the enriched iron-reducing consortia.

## AUTHOR INFORMATION

### Notes

The authors declare no competing financial interest.

## ACKNOWLEDGMENT

We thank Prof. Li Zhuang in Jinan University for her donation of *Geobacter sulfurreducens* PCA. This work was supported by the National Natural Science Foundation of China (91851202, 31970110, 51678163), Guangdong Provincial Science and Technology Project (2016A030306021, 2019B110205004), GDAS’ Special Project of Science and Technology Development (2019GDASYL-0301002), Guangdong technological innovation strategy of special funds (key areas of research and development program (2018B020205003), Open Project of State Key Laboratory of Applied Microbiology Southern China (SKLAM001-2018).

## REFERENCES

(1) Lloyd JR. 2003. Microbial reduction of metals and radionuclides. FEMS Microbiol Rev 27:411–425.

(2) Lovley DR, Anderson RT. 2000. Influence of dissimilatory metal reduction on fate of organic and metal contaminants in the subsurface. Hydrogeol J 8: 77–88.

(3) Byrne JM, Klueglein N, Pearce C, Rosso KM, Appel E, Kappler A. 2015. Redox cycling of Fe(II) and Fe(III) in magnetite by Fe-metabolizing bacteria. Science 347: 1473–1476.

(4) Yun J, Malvankar NS, Ueki T, Lovley DR. 2016. Functional environmental proteomics: elucidating the role of a c-type cytochrome abundant during uranium bioremediation. ISME J 10: 310–320.

(5) Logan BE, Rossi R, Ragab A, Saikaly PE. 2019. Electroactive microorganisms in bioelectrochemical systems. Nat Rev Microbiol 17: 307–319.

(6) Reguera G, McCarthy KD, Mehta T, Nicoll JS, Tuominen MT, Lovley DR. 2005. Extracellular electron transfer via microbial nanowires. Nature 435: 1098–1101.

(7) Yang Y, Xu M, Guo J, Sun G. 2012. Bacterial extracellular electron transfer in bioelectrochemical systems. Process Biochem 47: 1707–1714.

(8) Fortune WB, Mellon MG. 1938. Determination of Iron with o-Phenanthroline: A Spectrophotometric Study Ind Eng Chem Anal Ed 10: 60–64.

(9) Zhou S, Wen J, Chen J, Lu Q. 2015. Rapid measurement of microbial extracellular respiration ability using a high-throughput colorimetric assay. Environ Sci Tech Lett 2: 26–30.

(10) Xiao X, Liu Q, Li T, Zhang F, Li W, Zhou X, Xu M, Li Q, Yu H. 2017. A high-throughput dye-reducing photometric assay for evaluating microbial exoelectrogenic ability. Bioresour Technol 241: 743–749.

(11) Yang Z, Cheng Y, Zhang F, Li B, Mu Y, Li W, Yu H. 2016. Rapid Detection and enumeration of exoelectrogenic bacteria in lake sediments and a wastewater treatment plant using a coupled WO_3_ nanoclusters and most probable number method. Environ Sci Tech Lett 3: 133–137.

(12) Richter H, Lanthier M, Nevin KP, Lovley DR. 2007. Lack of electricity production by pelobacter carbinolicus indicates that the capacity for Fe(III) oxide reduction does not necessarily confer electron transfer ability to fuel cell anodes. Appl Environ Microbiol 73: 5347–5353.

(13) Luef B, Fakra SC, Csencsits R, Wrighton KC, Williams KH, Wilkins MJ, Downing KH, Long PE, Comolli LR, Banfield JF. 2013. Iron-reducing bacteria accumulate ferric oxyhydroxide nanoparticle aggregates that may support planktonic growth. ISME J 7: 338–350.

(14) O’Reilly SE, Watkins J, Furukawa Y. 2005. Secondary mineral formation associated with respiration of nontronite, NAu-1 by iron reducing bacteria. Geochem T 6: 67–76.

(15) Peretyazhko TS, Zachara JM, Kennedy DW, Fredrickson JK, Arey BW, McKinley JP, Wang CM, Dohnalkova AC, Xia Y. 2010. Ferrous phosphate surface precipitates resulting from the reduction of intragrain 6-line ferrihydrite by Shewanella oneidensis MR-1. Geochim Cosmochim Ac 74: 3751–3767.

(16) Qu ZJ, Li P, Zhang XX, Han KL. 2016. A turn-on fluorescent chemodosimeter based on detelluration for detecting ferrous iron (Fe^2+^) in living cells. J Mater Chem B 4: 887–892.

(17) Hirayama T, Okuda K, Nagasawa H. 2013. A highly selective turn-on fluorescent probe for iron (II) to visualize labile iron in living cells. Chem Sci 4: 1250–1256.

(18) Hirayama T, Tsuboi H, Niwa M, Miki A, Kadota S, Ikeshita Y, Okuda K, Hideko N. 2017. A universal fluorogenic switch for Fe(II) ion based on N-oxide chemistry permits the visualization of intracellular redox equilibrium shift towards labile iron in hypoxic tumor cells. Chem Sci 8: 4858–4866.

(19) Yang X, Wang Y, Liu R, Zhang Y, Tang J, Yang E, Zhang D, Zhao Y, Ye Y. 2019. A novel ICT-based two photon and NIR fluorescent probe for labile Fe^2+^ detection and cell imaging in living cells. Sensor Actuat B-Chem 288: 217–224.

(20) Ren J, Wu Z, Zhou Y, Li Y, Xu Z. 2011. Colorimetric fluoride sensor based on 1,8-naphthalimide derivatives. Dyes Pigments 91: 442–445.

(21) Chen X, Xu M, Wei J, Sun G. 2010. Two different electron transfer pathways may involve in azoreduction in Shewanella decolorationis S12. Appl Microbiol Biotechnol 86: 743–751.

(22) Xu M, Guo J, Kong X, Chen X, Sun G. 2007. Fe (III)-enhanced azo reduction by Shewanella decolorationis S12. Appl Microbiol Biotechnol 74: 1342–1349.

(23) Lu H, Deng T, Liu F, Wang Y, Yang X, Xu M. 2020. *Duganella lactea* sp. nov., *Duganella guangzhouensis* sp. nov., *Duganella flavida* sp. nov. and *Massilia rivuli* sp. nov., isolated from a subtropical stream in PR China and proposal to reclassify *Duganella ginsengisoli* as *Massilia ginsengisoli* comb. nov. Int J Syst Evol Microbiol doi: 10.1099/ijsem.0.004355.

(24) Li N, Than A, Sun C, Tian J, Chen J, Pu K, Dong X, Chen P. 2016. Monitoring dynamic cellular redox homeostasis using fluorescence-switchable graphene quantum dots. ACS Nano 10: 11475–11482.

(25) Ryckelynck N, Stecher HA, Reimers CE. 2005. Understanding the anodic mechanism of a seafloor fuel cell: interactions between geochemistry and microbial activity. Biogeochemistry 76: 113–139.

(26) Nicolaidou A, Nott JA. 1998. Metals in sediment, seagrass and gastropods near a nickel smelter in Greece: Possible interactions. Mar Pollut Bull 36: 360–365.

(27) Yang Y, Sun G, Guo J, Xu M. 2011. Differential biofilms characteristics of Shewanella decolorationis microbial fuel cells under open and closed circuit conditions. Bioresour Technol 102: 7093–7098.

(28) Zhao G, Li E, Li J, Liu F, Yang X, Xu M. 2019. Effects of flavin-goethite interaction on goethite reduction by Shewanella decolorationis S12. Front Microbiol 10:1623.

(29) Hori T, Aoyagi T, Itoh H, Narihiro T, Oikawa A, Suzuki K, Ogata A, Friedrich MW, Conrad R and Kamagata Y. 2015. Isolation of microorganisms involved in reduction of crystalline iron(III) oxides in natural environments. Front Microbiol 6:386.

(30) Wang Y, Ji Y, Wharfe ES, Meadows RS, March P, Goodacre R, Xu J, Huang WE. 2013. Raman activated cell ejection for isolation of single cells. Anal Chem 85: 10697–10701.

(31) Yang Y, Xiang Y, Sun G, Wu W, Xu M. 2015. Electron acceptor-dependent respiratory and physiological stratifications in biofilms. Environ Sci Technol 49: 196–202.

(32) Uma Vanitha M, Natarajan M, Sridhar H, Umamaheswari S. 2017. Microbial fuel cell characterisation and evaluation of Lysinibacillus macroides MFC02 electrigenic capability. World J Microbiol Biotechnol 33: 91.

(33) Azhar ATS, Nabila ATA, Nurshuhaila MS, Zaidi E, Azim MAM, Farhana SMS. 2016. Assessment and comparison of electrokinetic and electrokinetic-bioremediation techniques for mercury contaminated soil. IOP Conf Ser: Mat Sci Engin 160: 12077–12085.

(34) He H, Yuan S, Tong Z, Huang Y, Lin Z, Yu H. 2014. Characterization of a new electrochemically active bacterium, Lysinibacillus sphaericus D-8, isolated with a WO_3_ nanocluster probe. Process Biochem 49: 290–294.

(35) Ahmed B, Cao B, McLean JS, Ica T, Dohnalkova A, Istanbullu O, Paksoy A, Fredrickson JK, Beyenal H. 2012. Fe(III) Reduction and U(VI) Immobilization by Paenibacillus sp. Strain 300A, Isolated from Hanford 300A Subsurface Sediments. Appl Environ Microbiol 78: 8001–8009.

(36) Cao Y, Chen F, Li Y, Wei S, Wang G. 2015. Paenibacillus ferrarius sp. nov., isolated from iron mineral soil. Int J Syst Evol Microbiol 65: 165–170.

(37) Diez-Mendez A, Rivas R, Mateos PF, Martinez-Molina E, Julio Santin P, Antonio Sanchez-Rodriguez J, Velazquez E. 2017. Bacillus terrae sp. nov. isolated from Cistus ladanifer rhizosphere soil. Int J Syst Evol Microbiol 67: 1478–1481.

(38) Osman S, Satomi M, Venkateswaran K. 2006. Paenibacillus pasadenensis sp. nov. and Paenibacillus barengoltzii sp. nov., isolated from a spacecraft assembly facility. Int J Syst Evol Microbiol 56: 1509–1514.

